# Using methylation data to improve transcription factor binding prediction

**DOI:** 10.1101/2022.05.12.491666

**Authors:** Daniel Morgan, Dawn L. DeMeo, Kimberly Glass

## Abstract

Scanning the genome for the sequence patterns defined by Position Weight Matrices (PWM) is commonly used to estimate transcription factor (TF) binding “motif” locations. However, the PWM-based scores assigned to these inferred locations are associated with only modest accuracy when benchmarked against *in vivo* TF binding. One reason for this limited performance may be because PWMs do not incorporate information regarding the epigenetic context necessary for TF binding. To investigate this, we developed a framework to score inferred TF binding locations using CpG methylation data. We intersected motif locations identified using PWMs with methylation information captured in both whole genome bisulfite sequencing and Illumina EPIC array data for six cell lines, scored motif locations based on these data, and compared with experimental data characterizing TF binding (ChIP-seq). We found that, for most TFs, binding is better predicted using methylation-based scoring compared to standard PWM scores. In addition, our analysis shows that, while most TFs do not bind to methylated promoter regions, there are several exceptions to this rule, indicating that the role of methylation in TF binding may be cell-type and context specific. Finally, we also illustrate that our approach can be generalized to infer TF binding when methylation information is only proximally available, *i.e*. measured for nearby CpGs that do not directly overlap with a motif location. Overall, our approach provides a framework for inferring context specific TF binding using methylation data, a crucial initial step for understanding the impact of methylation on gene regulatory processes.

## Introduction

Modeling the regulatory mechanisms that determine cell fate, response to external perturbation, and disease state depends on measuring many factors, a task made more difficult by the plasticity of the epigenome. Transcription factor (TF) protein complexes binding to regulatory regions are one of the primary mechanisms that influence gene expression. DNA recognition sequences, summarized in position weight matrices (PWMs), can be compared with DNA sequence to identify potential TF binding sites across the genome. While PWM-based analyses have proven useful, they are purely *in silico* models thus limited in scope. Furthermore, since they are based on the DNA sequence, they cannot differentiate between intra-subject tissue or cell types.

Methylation is a widely studied epigenetic modification that reflects the dynamic regulatory changes the genome undergoes in response to the environment. Whole genome bisulfite sequencing (WGBS) data provides genome-wide information regarding methylation. In contrast, methylation arrays, such as the Illumina 450k and EPIC (850k) arrays, are high throughput assays that capture only a subset of methylation sites. However, these sites are selected, in part, based on their likelihood of involvement in gene regulation.

Several approaches have been developed to improve TF binding prediction by combining PWM-based motif predictions with epigenetic data such as chromatin accessibility^1, 2^, histone modifications^3, 4^, ATAC-sequencing^5^, and DNase hypersensitivity^6-8^. Previous studies using methylation data to predict TF binding have generally focused on specific cell line-TF combinations^9–13^. For example, Maurano et. al.^13^ specifically characterized CTCF binding affinity in the presence of methylation, but the study is not generalizable to other TFs. In fact, to our knowledge, such studies have never been carried out systematically across a broad range of TF and cell line combinations.

Methylation of gene regulatory regions (enhancers and promoters) is generally believed to inhibit TF binding. However, several studies have indicated a more nuanced relationship. For example, by combining aspects of cell-type specificity and scale, Lui et al.^14^ built and validated a random forest model to investigate how epigenetic changes impact TF binding^14^. In doing so, they identified a subclass of TFs that appeared to have an affinity for binding to methylated CpG sites. More recent genome-wide studies have also suggested that there are three classes of TF-binding in response to CpG methylation (affinity, restriction, and neutral)^15^. Similarly, Yin et. al.^16^ used SELEX profiling to systematically characterize TF binding preferences in the context of CpG methylation and classified TFs based on whether methylation had no effect, increased, decreased, or otherwise impacted their binding affinity^16^.

Ambrosini et. al.^17^ used SELEX profiling to analyze the accuracy of the PWM model in the context of ChIP-seq data^17^. This study determined that the PWM that was most predictive of a particular ChIP-seq experiment is generally not associated with the TF assigned to that PWM, but instead tends to be associated with another member of that TF’s family. This motivates the importance of improving upon PWM predictions using additional, context-specific genomic information. Here, we investigate improving the predictive capacity of TF-binding motifs by integrating sequence-based PWM predictions with methylation information. We compare with ChIP-Seq data to assess the predictive capacity of scoring motif locations based on the assumption that methylation inhibits TF binding.

In this study, we assess our approach using WGBS, methylation array, and TF ChIP-Seq data from six cell lines. This broad analysis allows us to gain a comprehensive understanding of the relationship between TF binding and methylation and to characterize the strengths and limitations of inferring TF binding using array versus sequence-based methylation information. We also use annotation information to distinguish the impact of methylation on TF binding in the context of various genomic elements. Finally, we present an option for generating predictions for TF binding when motif locations do not overlap with the methylation sites captured by a specific technology. As the scientific community continues to generate large amounts of methylation data, especially with the Illumina EPIC methylation array, we believe our approach and results will support the development of reliable and accurate models of gene regulatory mechanisms, including regulatory network inference. Because array methylation data is available in many large-scale human population studies, our approach could be applied to capture evidence for TF binding relevant to human disease.

## Methods

We used FIMO (Finding Individual Motif Occurrences)^18^ to scan the hg38 genome for 730 human TFs based on PWMs selected from CIS-BP^19^ and curated by the MEME suite (downloaded February 21, 2018). This analysis identified a list of genomic regions predicted to be bound by each TF (based on FIMO’s default cutoff of p<1×10^-4^); each genomic region has an associated PWM-score. For an individual TF the length of these genomic regions is constant; however, this length varies from 7 to 20bp depending on the TF. We analyze subsets of these genomic locations (also referred to as motif locations) by intersecting with other available genomic data using bedtools^20^. Our general approach is illustrated in **Figure 1**.

**Figure 1.**
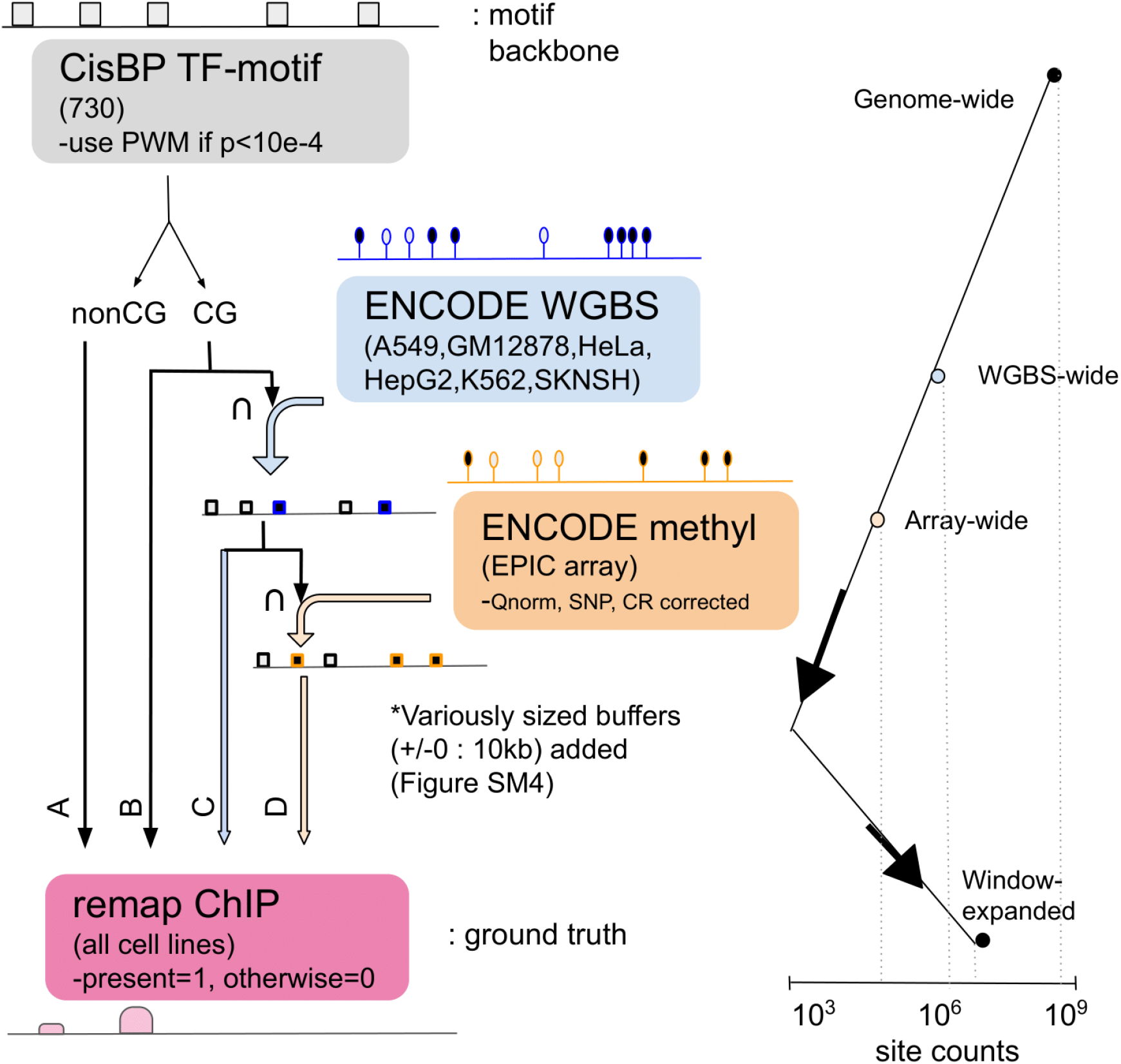
Intersection schema between data. Schematic workflow of how we intersected the data in our analysis. The intersection of all three data types with ChIP-seq data (CG motif ⋂ WGBS ⋂ array ~ ChIP) is the primary analysis presented in this manuscript and shown in **Figures 2-3**. This narrow view is then expanded by increasing the genomic region associated with each predicted transcription factor binding site (**Figure 4**).

Next, we downloaded ChIP-seq data for six cell lines (A549, GM12878, HeLa, HepG2, K562, SKNSH) from ReMap^21^. For each PWM, we compared its identified motif locations to cell line specific ChIP-seq binding locations for the PWM’s assigned TF. For a given TF and cell line, this resulted in two values associated with each motif location: a score from FIMO based on the PWM match and a binary (0/1) score based on whether a ChIP-seq peak overlapped with that location in a specific cell line. We then used the binary ChIP-seq score as a gold standard to calculate TF and cell line specific AUROC (Area Under the Receiver-Operator Characteristic curve) scores. Globally, more than 9% of motif locations overlapped with a ChIP-seq peak; across the tested TF cell line combinations the median percentage of motif locations with a ChIP-seq peak was 5%.

Since methylation generally occurs at CpG dinucleotides, we parsed TF motif locations into two mutually exclusive groups, the first consisting of locations not containing CpG sites and the second consisting of locations containing CpG sites. We scored motif locations based on the PWM-score and calculated the AUROC separately using each set of motif locations. We observe no significant differences, indicating that the TF locations which we can score using methylation data (which, by definition, will contain a CpG) is not a biased subset (**Figure S1**).

Next, we downloaded WGBS and EPIC array methylation data for six cell lines (A549, GM12878, HeLa, HepG2, K562, SKNSH) from ENCODE^22^ (accessed Jan 15, 2020). We determined the correlation between these two methylation technologies and observed technical artifacts for low read depths (**Figure S2**). Based on this analysis, for each cell line we filtered to include only methylation sites with a WGBS read depth ≥10; this allowed us to maximize correlation between the technologies while minimizing loss of sites. We then used these data to score TF motif locations that contain at least one methylation site measured in both the WGBS data (among sites with a read depth ≥10) and in the array methylation data. We assumed that methylation inhibits TF binding, and thus for each motif location we calculated a score (*m*) that is inversely related to the methylation measure.

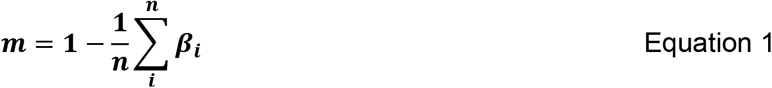

Based on **Equation 1**, when multiple methylated CpGs (mCpGs) are present within the same motif location, we take their average, although we also explored alternative ways of scoring motif locations that overlap multiple methylation sites, including taking the median, min, or max (**Figure S3**). In **Equation 1** *β_i_* represents the methylation value for any CpG site *i* that overlaps with the motif location. To derive *β_i_* for the WGBS data, we divide the reported methylation value by 100 to ensure a range of 0-1 to be consistent with the range of the beta values associated with the array technology. In this way we upweight hypomethylated motif regions (*m* approaches 1) and down-weight hypermethylated regions (*m* approaches 0). This process results in cell-line specific scores for a subset of motif locations -- those that overlap with CpGs with at least 10 reads in WGBS that are also included in the array data. This greatly limits the number of motif locations being assessed, primarily due to the array data only measuring a subset of all possible methylation sites (**Figure 1D, Table 1**). However, it also enables us to directly compare the results between WGBS and the more widely available EPIC array data.

**Table 1.**
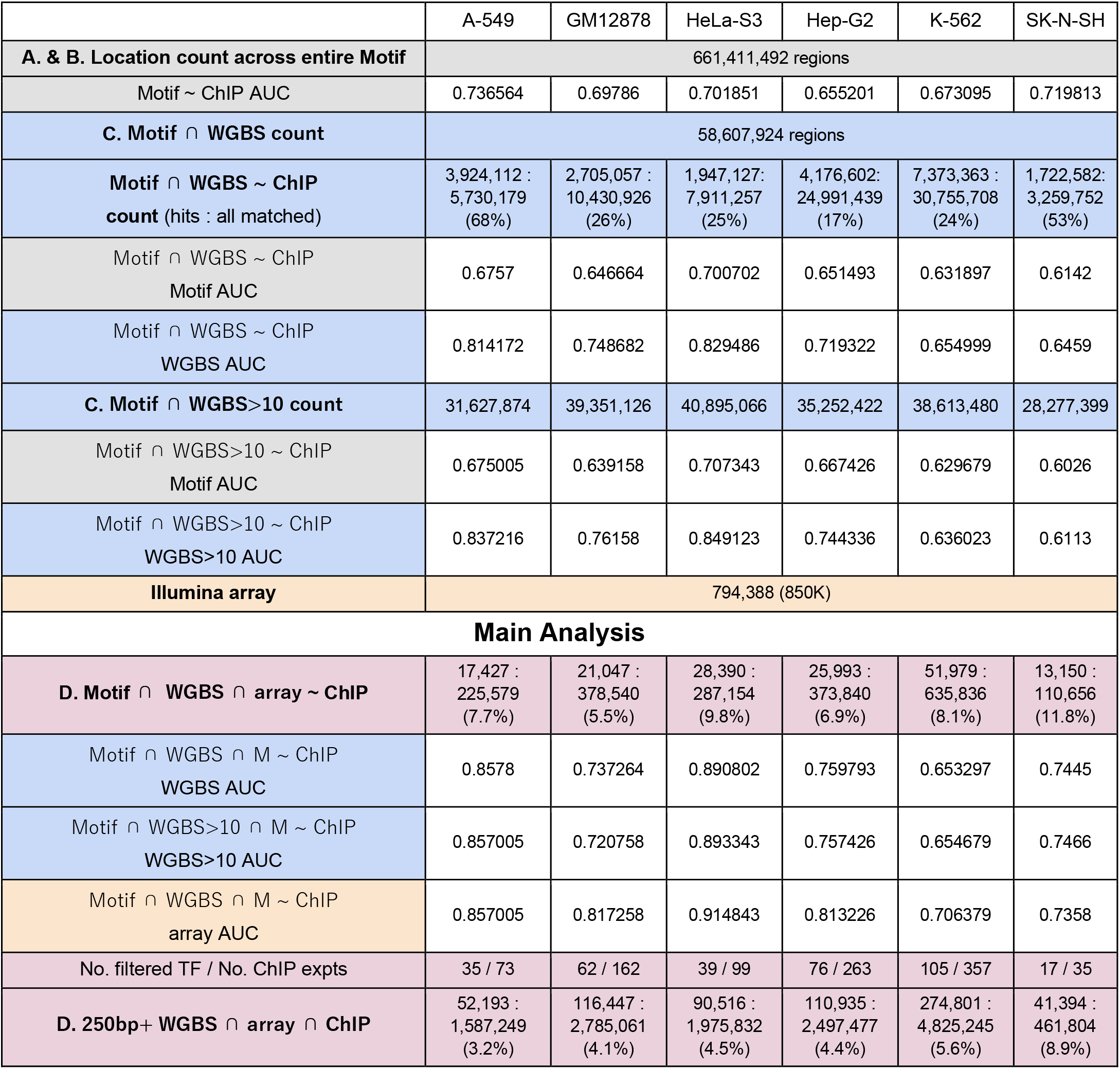
Counts of intersecting regions across omics. While generally on the same order of magnitude, all cell lines have a slightly different number of assessed regions across the various omics. Intersecting the regions available for each of the omics determines the number of transcription factors and binding sites assessed in our analyses. White elements in the table display the median AUC across all TFs in a cell line while colored elements show the number of elements (TFs or regions) being assessed. The first column of this table references **Figure 1**.

We use this reduced set of TF motif locations together with EPIC array annotation to investigate if genomic context impacts the inhibitory role of methylation on TF binding. We note that EPIC array annotations generally hold up well against external annotation tools^23, 24^. In the annotation analysis, we determine the primary annotation of each methylation site, and then score motifs by averaging across CpGs that have a specific associated primary annotation (*A*):

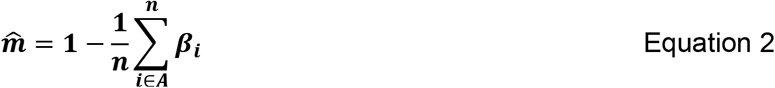

This means that each methylation site is considered only once within the annotation-based AUROC analysis.

Finally, to put our observations into the context of the current literature, we extracted the results of a study^16^ that used data from the SELEX platform to characterize the effect of methylation on the binding of human TFs. From this paper, we focused on the 292 TFs categorized by the authors as either MethylPlus (175) or MethylMinus (117), and disregarded TFs labeled as having “multiple effects”, “Little Effect”, or “no CpG”. Of these 292 TFs, 55 were also evaluated in our motif scan and had ChIP-seq data in at least one cell line.

## Results

### Predicting TF binding using methylation data

We leveraged four data types in our analysis: (1) genome-wide motif locations derived by running FIMO on PWMs from CIS-BP and curated by MEME, (2) WGBS and (3) EPIC array methylation for six cell lines from ENCODE, and (4) ChIP-seq peaks for these same cell lines from ReMap. We observed no statistically significant differences in predicting TF binding between motif locations containing CpGs and those devoid of CpGs (**Figure S1**). Furthermore, we find that the methylation levels of CpGs captured by both the array and WGBS technologies are highly correlated, although we observe a lack of resolution for low-coverage CpG sites in the WGBS (**Figure S2**). Thus, we thresholded to only include CpGs from WGBS data for which the sequence reads depth was greater than or equal to 10. We note that this does not significantly reduce the number of methylation sites considered in our main analysis since most sites (>75%) that are measured by the Illumina array have a read depth greater than or equal to 10 in the WGBS data.

Next, for each of the six cell lines, we scored motif locations that overlapped at least one methylation site that is assayed by both the WGBS and array technologies. We compared methods of aggregating methylation information when multiple methylation sites overlap with the same motif location. We observed the scores based on different aggregative approaches to be highly correlated (**Figure S3**). Based on this analysis, when multiple methylation sites are associated with the same motif location, that location is given a single value based on averaging the methylation levels of all associated sites (**Equation 1**). This process resulted in two cell-line specific scores for each characterized motif location: one based on methylation values from WGBS and one based on methylation values from the EPIC Illumina array.

For each TF motif and cell line, we compared these scores with ChIP-seq data to quantify how accurately they predict TF binding. The distribution of the results, as quantified using the AUROC, is shown in **Figure 2A**. TF specific AUROCs and other comparisons can be found in **Figure S4**. We observe a low level of prediction when scoring motifs locations based on their original PWM score (AUROC=0.55-0.65) consistent with previous findings^25^. However, when scoring motif locations based on methylation data we observe much higher accuracy, with an overall AUROC increase of 0.2 (**Figure 2A**) and individual transcription factors increases of 0.1-0.4 (**Figure S4**). These results strongly suggest that methylation data can be used to predict TF binding when ChIP-seq data is unavailable. These results also confirmed our previous assumption that DNA methylation should generally inhibits TF-binding (**Equation 1**). While both WGBS and array methylation information improves our ability to predict TF binding compared to PWM-scores, the extent of improvement is significantly greater for the Illumina array data compared to the WGBS (p-value 1×10^-36^ based on a paired t-test). This may be due to the fact that the methylation values obtained using sequencing technology are dependent on the coverage of individual sites, while the values obtained using arrays are continuous and thus associated with a more fine-grained assessment of a site’s methylation level.

**Figure 2.**
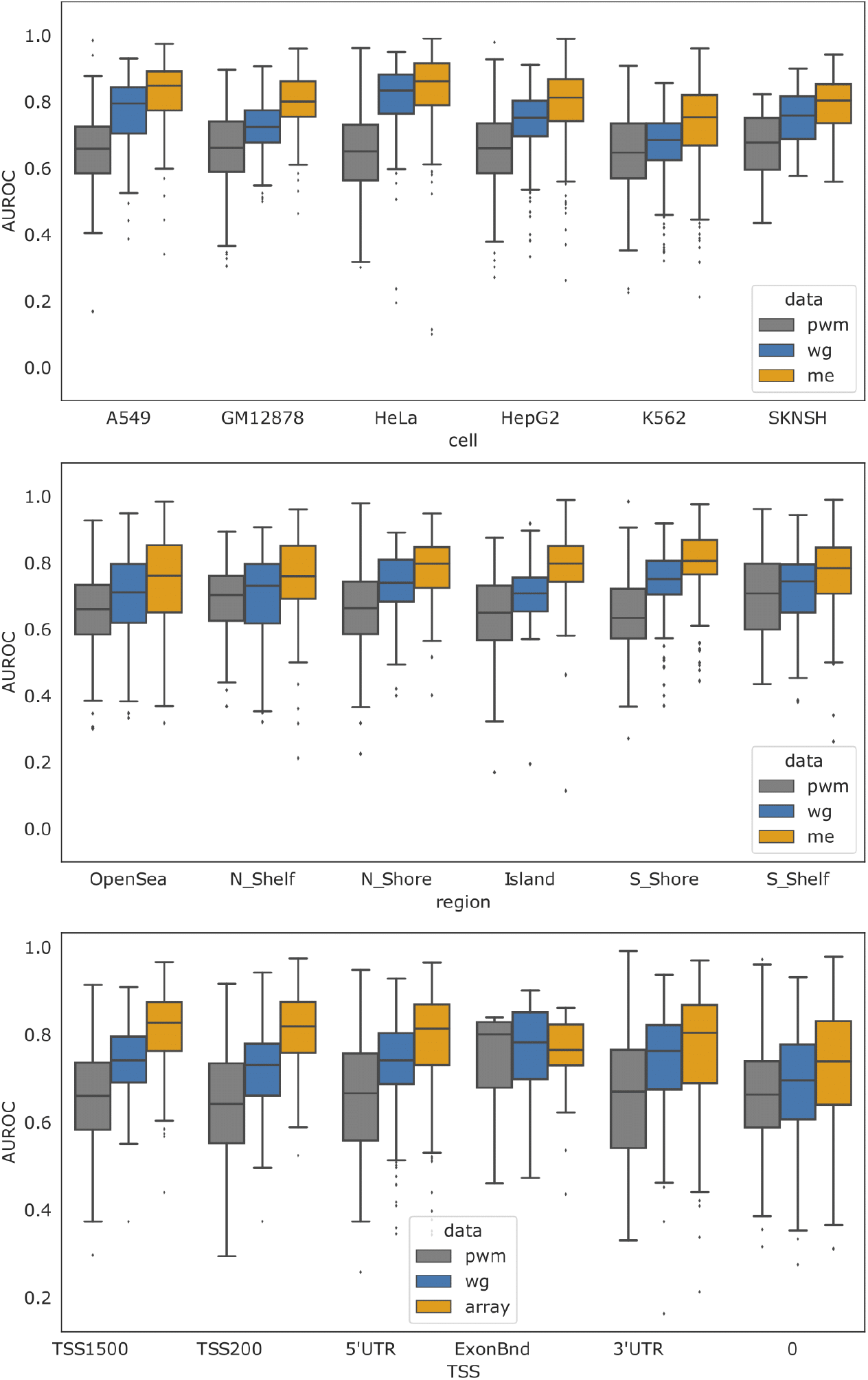
AUROC comparisons across cell lines and genomic annotations. **(A)** Boxplots representing the distribution of TF AUROC scores for each cell line. All AUROCs improve using methylation-based scoring compared to PWM-based scoring at a significance level of p<2×10^-3^. AUROC scores for individual TFs are shown in **Supplemental Figure S4**. **(B)** Boxplots representing the distribution of TF AUROC values when assessed within different genomic regions, indicating that the overall improvement in predicting TF-binding can be attributed to region-specific improvement. **Supplemental Figure S5** has TF specific breakdowns. **(C)** Boxplots representing the distribution of TF AUROC values when assessed in the context of gene regulatory regions, indicating that the overall improvement in predicting TF-binding can be attributed to improvements near the TSS. **Supplemental Figure S6** has TF specific breakdowns. Note that the ExonBnd has an order of magnitude less associated motif locations. 0 refers to unlabeled. pwm = PWM; me=methyl array; wg=WGBS.

Next, we used Illumina methylation array annotations to evaluate if the methylation information associated with different genomic contexts play distinct roles in predicting TF binding. To do this, we calculated context-specific methylation scores for each motif location (**Equation 2**); these scores are identical to the previous methylation scores except in the rare case where multiple methylation sites annotated to different genomic contexts overlap with the same motif location (see **Methods**). For each TF and cell line, we selected the subset of motif locations associated with a particular annotation and benchmarked these scores by comparing with ChIP-seq; this resulted in three AUROC values (for the PWM score, WGBS score, and array methylation score) for each TF, cell line, and annotation combination. We then averaged AUROC values across TFs assessed in multiple cell lines, resulting in one value per TF for each annotation. The distribution of the results for annotations related to gene body proximity and TSS region are shown in **Figure 2B-C**. TF specific results and other comparisons can be found in **Figure S5-S6**.

This analysis indicates that methylation information associated with promoter regions increases predictive performance, with the highest AUROC improvement occurring when assessing motif locations within CpG islands, followed by the shores and the shelves. Only marginal improvement is observed for motif locations annotated to open sea regions (**Figure 2B**). In addition, methylation sites annotated to regions near the transcriptional start site (TSS) appear to greatly enhance our ability to predict TF binding, with the highest AUROC increase observed when using motif locations annotated to promoter regions (**Figure 2C**). In contrast, methylation sites annotated to exons had a predictive capacity no better than the original PWM score.

### Context-specific patterns in TF binding predictions

Our analysis thus far indicates that methylation information can be used to predict TF binding. However, given the plasticity of the epigenome, we next investigated if predictive performance varies for individual TFs across different cell lines and genomic contexts. To do this, for each TF and within each cell line, we scored and determined the predictive capacity of motif locations annotated to promoters versus the gene body and compared to the predictive capacity of the PWM-score (**Figure 3A**). We find that for most, but not all, individual TFs and cell lines, TF binding is better predicted when scoring motif locations using methylation information (**Figure 3A** above diagonal), although for each scoring scheme and context there are a small number of poor predictions (AUROC values around 0.5). We also note that the AUROC values are significantly higher when evaluating motif locations associated with promoter regions (TSS200 and TSS2500) as compared to the gene body (3’ UTR, 5’ UTR and Exon Boundary) when using the array data (two-sided t-test p-value = 0.002). However, this is not the case for WGBS data (two-sided t-test p-value = 0.177). This further supports our previous observation that promoter specific methylation, especially that captured using array technology, is enhancing our ability to correctly predict TF-binding (see **Figure 2C**).

**Figure 3.**
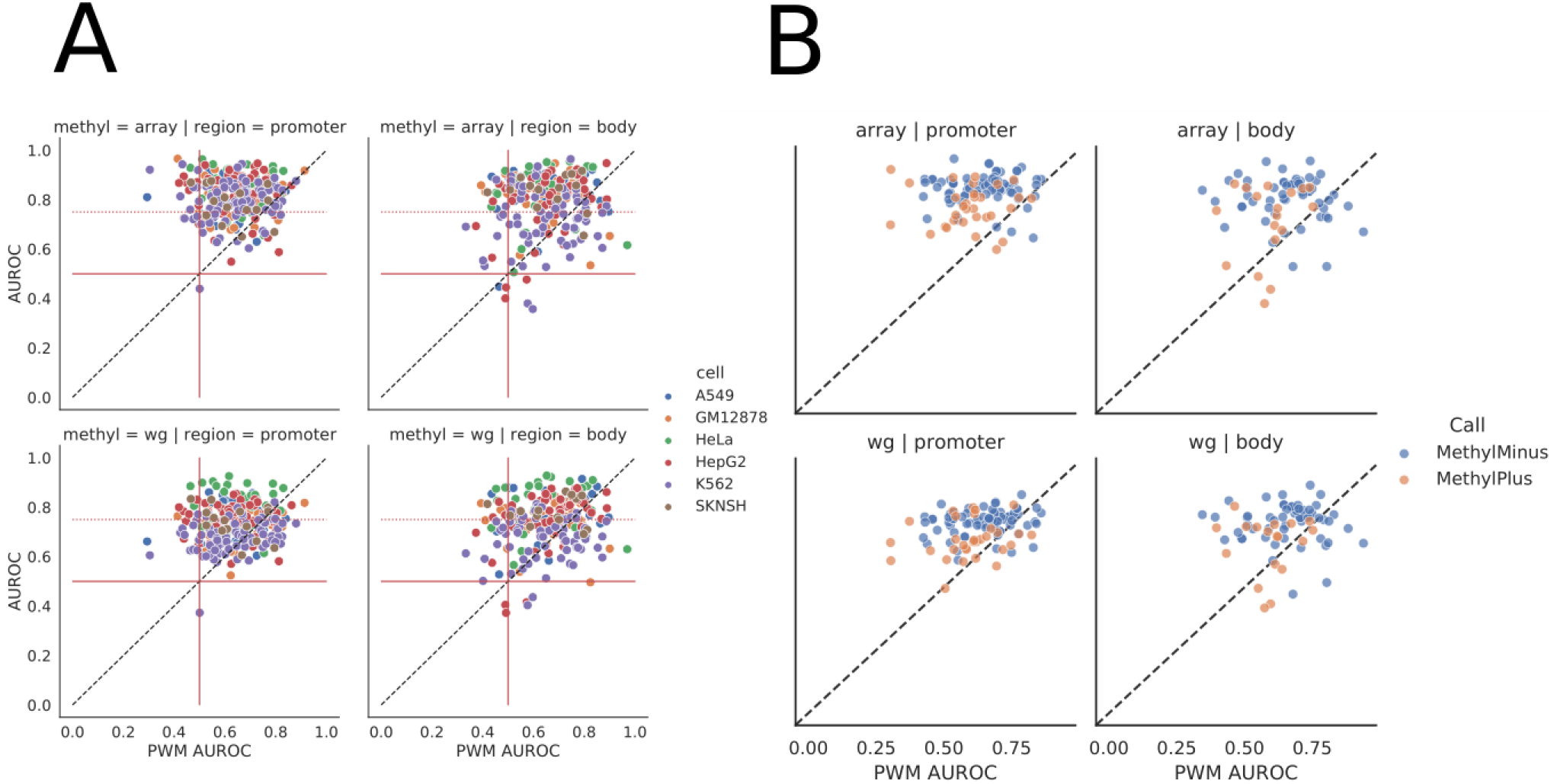
Performance of individual TFs in different genomic regions. **(A)** Performance of PWM-based scoring for each TF and cell line (each point is a unique TF/cell-line combination), compared to scoring based on methylation type (WGBS or methylation array) and genomic context (gene body or promoter). Solid red lines at 0.5 are displayed to highlight TFs with little signal in either methylation type. A dotted red line at 0.75 is plotted along the methylation-specific y-axis to distinguish minor from major signal. Several CEBP family members lie below 0.5 AUROC and are listed in **Supplemental Figure S6F**. **(B).** Performance of PWM-based scoring for each TF (each point is a unique TF), compared to the performance based on methylation type (WGBS or methylation array) and genomic context (gene body or promoter). The color of the circles indicates whether the TF belongs to the MethylMinus (blue) or MethylPlus (yellow) class as defined in Yin et al. In contrast to (A), in these plots each point is a unique TF averaged across cell lines to match cell-line agnostic Yin labels.

We also observe that, primarily in the context of the gene body, the binding of some TFs is better predicted using the original PWM score rather than the methylation-based scores (**Figure 3A** below diagonal). In particular, there are several TFs, mostly belonging to the CEBP family (**Figure S6F**), for which the presence of DNA methylation appears to enhance, rather than inhibit TF binding in a specific context (AUROC values of about 0.4). However, binding of these TFs is also generally poorly predicted by the PWM score – all but one has an AUROC based on the PWM score of less than 0.6, with half having a PWM-based AUROC of approximately 0.5 or less. Therefore, it is difficult to draw a specific conclusion for these TFs. However, we note that these results are consistent with previous observations that some CEBP proteins can bind to both unmethylated and methylated versions of their primary DNA recognition sequence, and that methylation may enable them to bind to additional, non-canonical binding sites^26–28^.

Next, we compared our results to the “MethylMinus” and “MethylPlus” transcription factor classes identified in Yin et. al.^16^, which relied upon protein binding SELEX microarray data to determine transcription factor affinity to bind to methylated DNA. To start, similar to the approach we took in **Figure 2B-C**, we averaged AUROC values for TFs assessed in multiple cell lines, resulting in one value for each TF. Then, for the subset of TFs with a designated class in Yin et. al. **(~43%)**, we plot these values compared to AUROCs based on the PWM-score (**Figure 3B**). We also compared these values between the TFs that Yin et. al. designated as MethylPlus from MethylMinus and find that both array and WGBS derived scores can distinguish these classes within promoters (p-values equal to 4.89×10^-7^ and 1.14×10^-7^, respectively, based on paired t-test) and, to a lesser extent, the gene body (p-values 5.79×10^-4^ and 1.24×10^-4^).

### Leveraging methylation information from proximal CpGs

Most TF motif locations do not directly overlap with a CpG site covered in the array data and therefore were excluded from our primary analysis. For example, of more than 661 million genomic locations identified in our original motif scan, only 31.6 million overlapped with a CpG methylation site covered in the A549 WGBS with a read depth of more than 10; this shrunk to only 225.6 thousand locations when we restricted to methylation sites contained in the array data and motif locations that corresponded to potential transcription factor binding sites for which we had ChIP-seq data (**Table 1**). Given this limited coverage, we next investigated our ability to predict TF binding when using methylation values from CpGs that are nearby, but do not necessary directly overlap with, a given motif location. To do this, we added a window before and after each identified motif location, essentially enlarging the genomic region covered by each. Next, we intersected methylation sites from WGBS and array data with motif locations that had been expanded based on various window sizes, and scored these locations based on **Equation 1**. We investigated adding windows of +/-5, 10, 20, 50, 100, 250, 500, 1000, 2500, or 5000 bp. These window sizes allow us to investigate the scale at which CpG methylation is predictive of TF binding.

**Figure 4** shows the distribution of AUROC values across all TFs and cell lines when using the PWM score, the baseline methylation score when using no window (equivalent to the results from **Figure 2A**), and the various methylation scores obtained when expanding motif locations based on each of the window sizes. We find that the distribution of AUROC values is highly stable up to a window size of +/-100bp. This suggests there is local correlation in methylation levels across short ranges of the genome, consistent with what has been described in the literature^11^. Interestingly, as the window size increases, the improvement of the methylation score compared to the PWM score becomes more consistent between the WGBS and methylation array data (**Figure S7**). This may be due to the fact that averaging over multiple sites increases the resolution of the WGBS-based methylation score, making it more comparable to the array-based methylation score.

**Figure 4.**
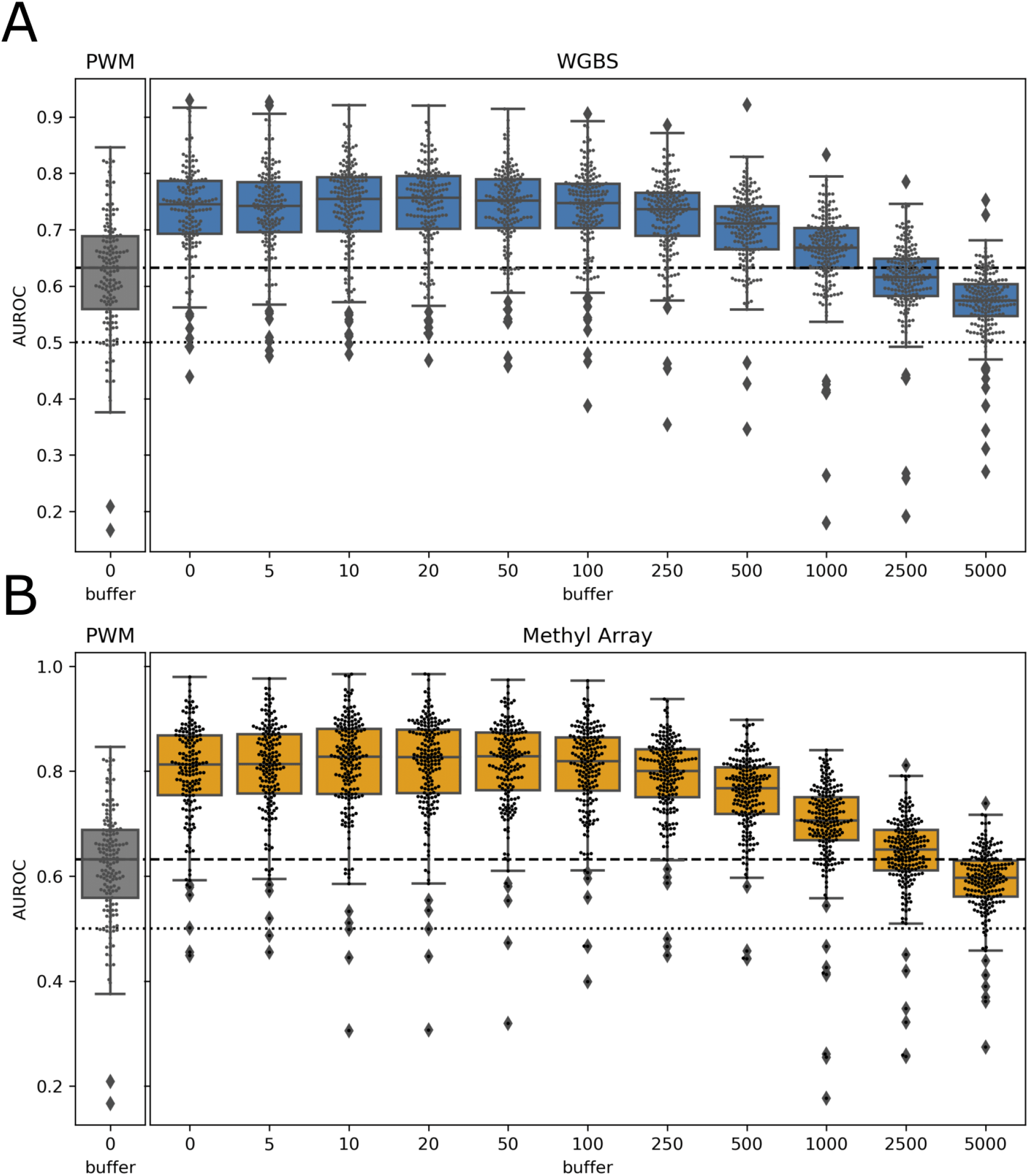
Performance when using nearby CpGs to score motif locations: Boxplots representing the distribution of all TF AUROC scores in all cell lines when the methylation-based score is derived from nearby CpGs. The x-axis indicates the window (or buffer) around the CpG sites that was used. A buffer of 0 indicates direct overlap of the CpG site with the TF motif, as was done for the analyses shown in Figure 3-4. We observe similar performance up to a window size of about +/-100bp, after which there is a sharp drop-off in predictive performance.

Overall, this analysis indicates that we can use methylation information to predict TF binding for motif locations that do not directly overlap with a CpG site assayed by a specific technology. Instead, methylation sites that are within +/-100 bp of a motif location can be used to score that location and predict TF binding.

## Discussion

In this manuscript we evaluate using methylation data to predict *in vivo* transcription factor binding. To do this, we scored predicted transcription factor binding sites using methylation levels for CpGs captured using both whole genome bisulfite sequencing and on Illumina 450K microarrays across six cell lines; we then benchmarked by comparing with ChIP-seq data for transcription factors from those same cell lines. We found that our methylation-based score consistently performed better at predicting transcription factor binding compared to the original PWM score.

We also observed that our methylation-based scoring approach was most predictive of *in vivo* transcription factor binding when the CpGs used to score binding sites were annotated to gene regulatory regions, including CpG islands, shelves, and shores, as well as promoters. We note that this context-specific analysis is limited by our use of annotations from the Illumina methylation array. However, our results are consistent with the current understanding of the regulatory role of DNA methylation, namely, that it can inhibit transcription factor binding in gene regulatory regions, thereby repressing the associated genes’ expression. Furthermore, the wide use of the Illumina platform allows us to interpret these findings in light of other studies in the field.

We also note that for a small number of transcription factors, our methylation-based score had minimal predictive capacity (**Figure S6F**). Interestingly, many of these transcription factors are CEBPs. CEBPβ has previously been found to bind nearly equally well to both methylation and unmethylated versions of its binding site^27^, and both CEBPα and CEBPβ have been shown to bind to a methylated version of the consensus CREB binding site^26^, indicating that methylation levels may have only minimal impact on CEBP binding to the DNA. Thus, given the robust, redundant, and highly plastic nature of methylation, our results suggest that methylation is not universally inhibitory. However, we note that given the relative few instances in which we observe this behavior, it is impossible to conclude with certainty whether these results are biological or a result of noise or platform error.

Following this observation, we also investigated whether our results were consistent with previously observed transcription factor binding behavior in the context of methylation, as reported in (Yin *et. al*. 2017)^16^; this paper used SELEX technology to determine if methylation of CpGs within transcription factor motif(s) tended to decrease, increase, or had little/no effect on the overall ability of transcription factors to bind to that sequence. We found only minimal distinction in our ability to predict binding for transcription factors associated with the TF classes defined in this paper, although there was a small, but statistically significant, increase in the associated AUC values for the set of transcription factors the paper identified as having a decreased ability to bind in the presence of CpG methylation. Differences in platform technology and overall analysis approach likely contribute to the minimal distinction between the transcription factors associated with these class categorizations in our analysis.

Finally, we note that, by focusing on CpGs captured using two different methylation technologies, in our main analysis we were only able to assign methylation scores for a small subset of all potential transcription factor binding sites. However, we also evaluated using a methylation score that incorporated methylation levels from CpGs that are nearby, but not necessarily overlapping with, a predicted binding site. We observed consistent predictive performance when using methylation scores that incorporate information for CpGs that are up to 100bp away from predicted transcription factor binding sites. By considering proximal CpGs we can score over ten times more transcription factor binding sites compared to only using direct overlap; this includes approximately one third of the transcription factor binding sites in gene promoter regions, areas that typically have high coverage on methylation microarrays.

Together with our observation that the highest predictive performance is for transcription factor binding sites located in gene regulatory regions, our analysis sets the stage for using high-throughput methylation arrays to estimate gene regulatory networks. The large amount of methylation data that has already been generated for longitudinal and disease cohorts means that we may now be able to more fully characterize biological systems and their evolution across disease progression, aging, etc., something that would be impossible using only PWM scores to predict transcription factor regulatory networks. This is an important future direction for our group^25^.

The main strength of our analysis is that it provides a high-level, comprehensive assessment of how methylation data can be used to provide condition-specific prediction of *in vivo* transcription factor binding. This has profound implications for the type of information that can be extrapolated from the large amount of methylation data that already exists and is being generated. However, our approach is also limited. First, we focused on methylation data from cell lines. Although this allowed us to systematically benchmark our results using a large amount of ChIP-seq data, methylation data in cell lines likely has distinct differences from that of primary cells and that gathered for complex diseases or from tissue samples. Secondly, we focused on transcription factors for which there is a known PWM and for which we had ChIP-seq data; this may be a slightly biased subset. Finally, we focused our assessment on transcription factor binding sites that directly overlapped with CpGs assayed in both whole genome bisulfite sequencing data and methylation array data. This allowed is to directly compare these technologies, but it also severely restricted our search space. However, we also performed an analysis that indicated that we could score transcription factor motif locations that do not directly overlap with an assayed CpG, greatly expanding the potential applicability of our approach. Together with our observation that methylation associated with promoter regions and CpG islands increased our predictive performance and the fact that methylation arrays are generally enriched for gene regulatory regions, we believe our approach could be applied to understand gene regulation or to model gene regulatory networks.

In summary, in this manuscript we investigate the use of scoring predicted transcription factor binding sites using methylation data. We show that methylation-based scores significantly improved our ability to predict experimental (ChIP-seq) transcription factor binding compared to the original PWM-based score. Importantly, our results are robust across transcription factors, cell lines, and methylation platforms. We believe using methylation data to improve transcription factor binding prediction provides a framework that can be used to better understand the mechanistic and regulatory drivers of biological systems.

## Acknowledgments

T32HL007427, P01HL132825, P01HL114501, R01HL152735, R21HL156122, K25HL133599, R01HL155749

## Code Access

https://github.com/dcolinmorgan/mili_benchmark/blob/master/notebook/v8_channing_methyl_benchmark.ipynb

## Data Access

ChIP-seq from ReMap2020 Homo sapiens -- all peaks: https://remap.univ-amu.fr/download_page

WGBS from ENCODE: https://www.encodeproject.org/matrix/?type=Experiment&status=released&assay_slims=DNA+methylation&biosample_ontology.classification=cell+line&assay_title=WGBS&biosample_ontology.term_name=A549&biosample_ontology.term_name=K562&biosample_ontology.term_name=GM12878&biosample_ontology.term_name=HeLa-S3&biosample_ontology.term_name=HepG2&biosample_ontology.term_name=SK-N-SH

meArray from ENCODE https://www.encodeproject.org/matrix/?type=Experiment&status=released&award.project=ENCODE&files.platform.term_name=Illumina+Infinium+Methylation+EPIC+BeadChip&biosample_ontology.term_name=A549&biosample_ontology.term_name=K562&biosample_ontology.term_name=GM12878&biosample_ontology.term_name=HeLa-S3&biosample_ontology.term_name=HepG2&biosample_ontology.term_name=SK-N-SH&assay_title=DNAme+array

## Supplemental Figures

**Supplemental Figure S1.**
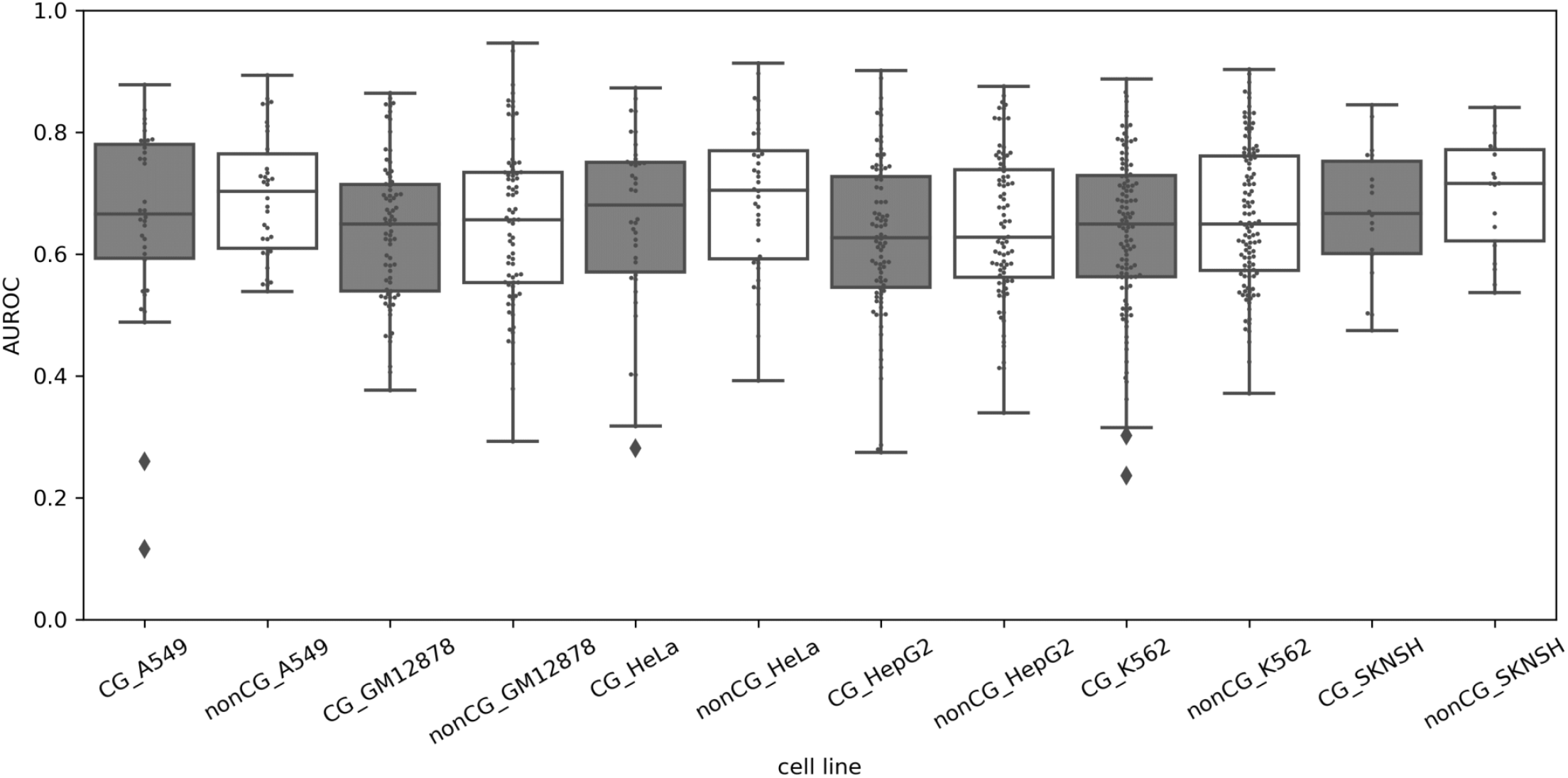
Comparing predictions between motif locations with or without a CpG. A comparison of the PWM score’s ability to predict *in vivo* TF binding (ChIP-seq) when limited to predicted binding sites that contain, or do not contain, a CpG (see **Figure 1 A&B**). A T-Test comparing performance within each of the cell lines confirms no significant difference. P-value for A549: 0.233128, GM12878: 0.566850, HeLa: 0.235387, HepG2: 0.336910, K562: 0.127878, SKNSH: 0.365327.

**Supplemental Figure S2.**
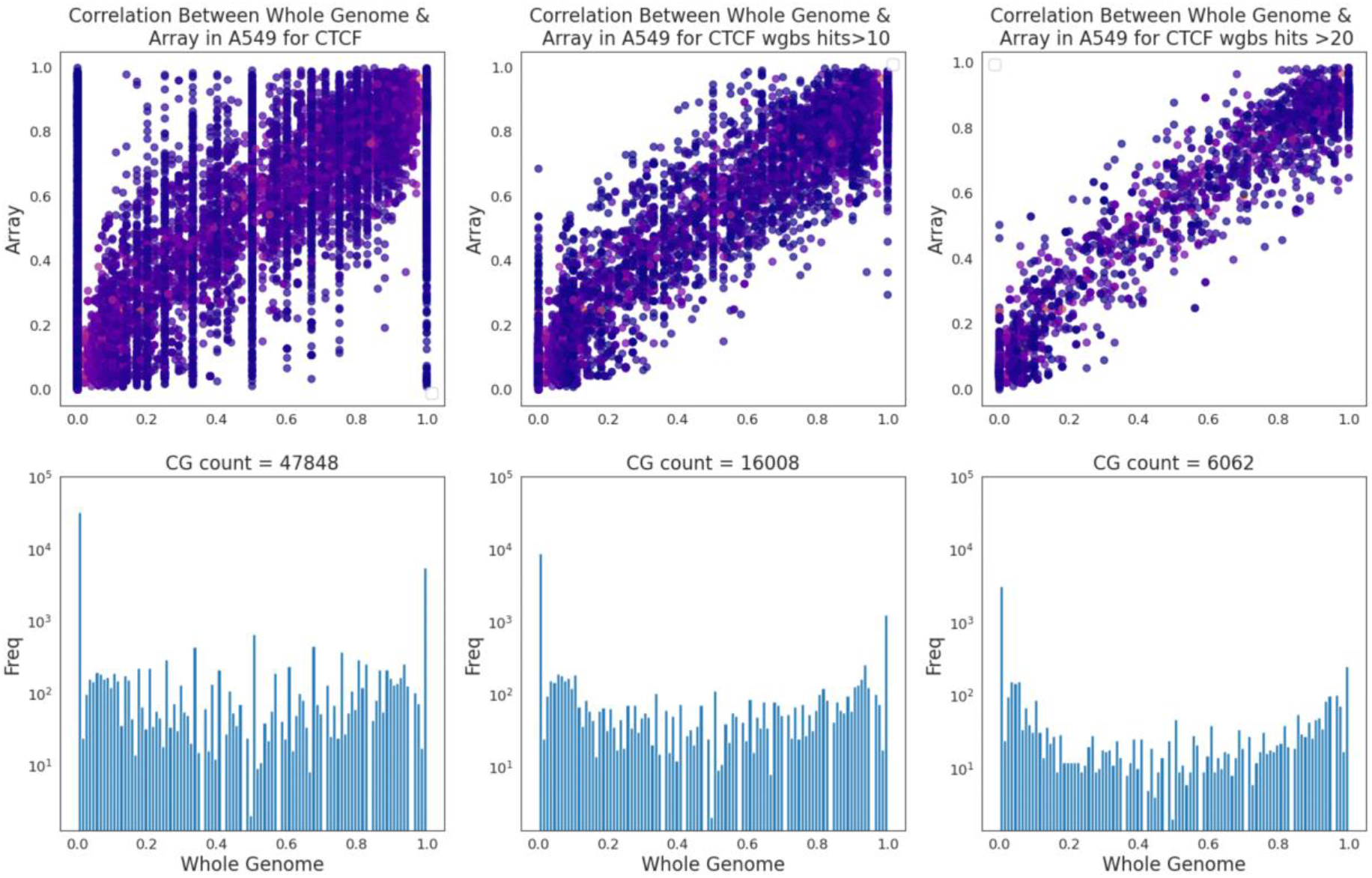
WGBS sequence depth analysis for a representative example: CTCF in A549. (**A**) Scatter plots comparing WGBS to methylation array values when restricting to sites in the WGBS with a read depth greater than 0, 10, or 20. **(B)** Histograms of WGBS values when the sites are restricted to those with a read depth greater than 0, 10, or 20.

**Supplemental Figure S3.**
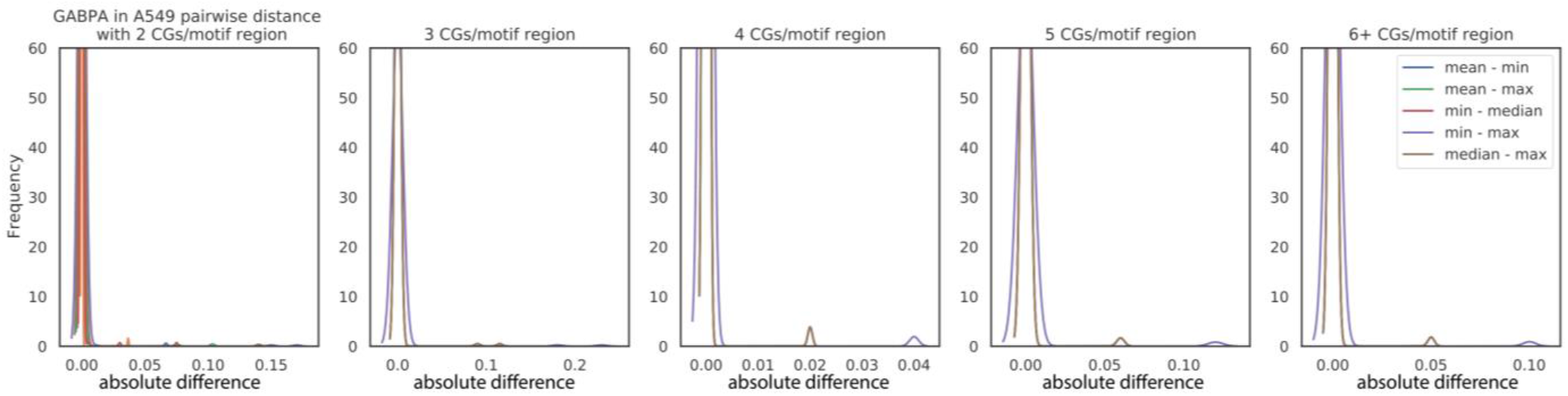
Absolute pairwise difference when using different measures to summarize methylation values when multiple CpGs overlap with a PWM, using CTCF in A549 as an example. Density curves (via fit kernel density estimates) comparing various methods for aggregating CpG values, broken into groups depending on the number of CpGs found within the predicted transcription factor binding site (motif hit): 2, 3, 4, 5, or 6+ CpGs per motif. Note that the mean-median comparison was too narrow to visualize.

**Supplemental Figure S4.**
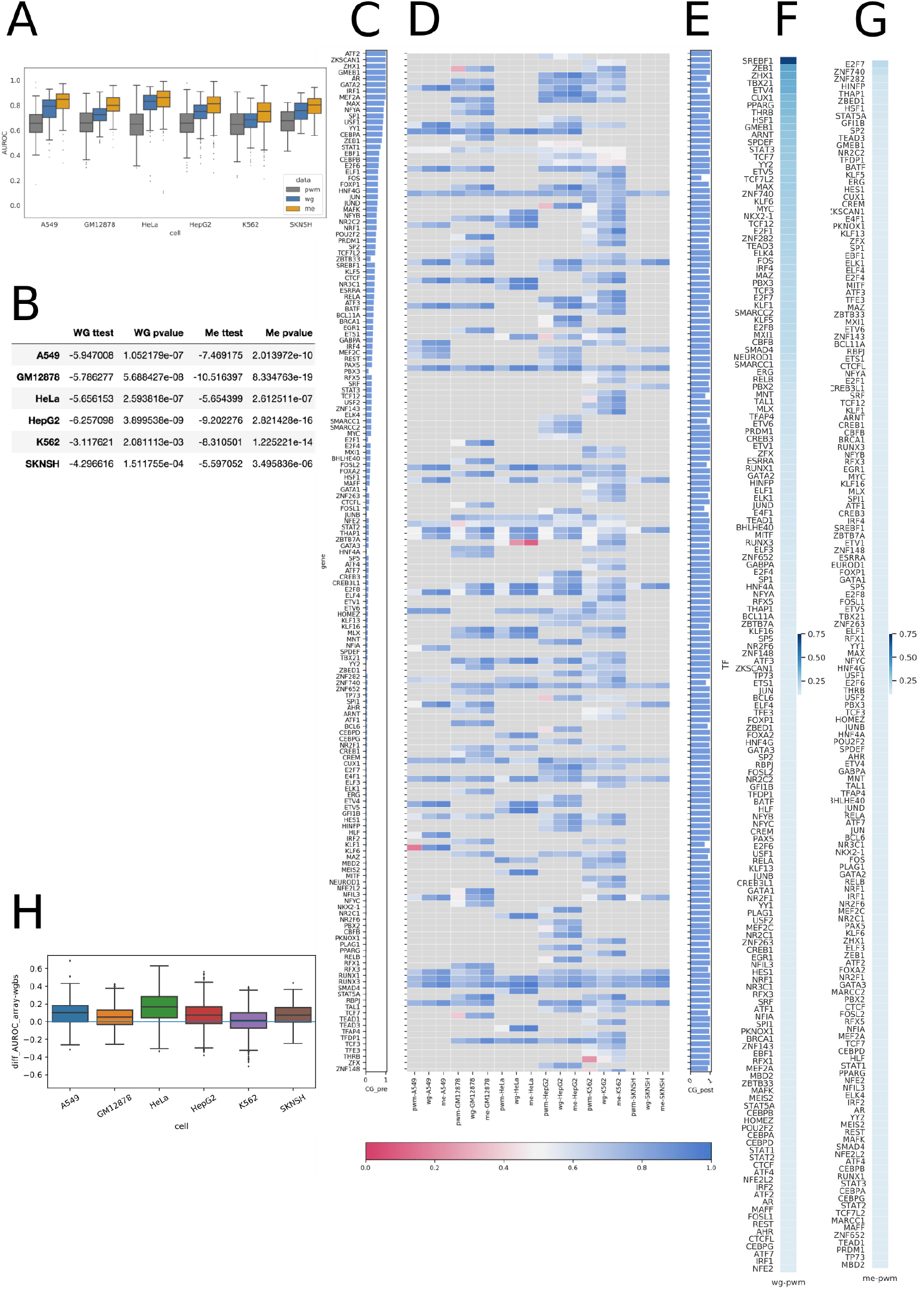
Details supporting main analysis. **(A)** Same plot as shown in **Figure 2A.** Each point is a TF-cell line combination. **(B)** Table showing the T-test statistic and p-value when comparing the AUROC distributions for the score to WGBS and methyl array. **(C)** Bar chart showing the percentage of motif locations containing a CpG before intersecting with the methylation data. **(D)** Heatmap of individual cell line specific TF performance. **(E)** Bar chart showing the percentage of motif locations containing a CpG after intersecting with the WGBS and array data demonstrating this process is selecting for CpG containing motif locations. **(F)** Difference in AUROC performance for each TF (averaged across cell-lines) comparing WGBS minus PWM-based scoring. **(G)** Difference in AUROC performance for each TF (averaged across cell lines) comparing Methyl Array minus PWM-based scoring. **(H)** Distribution of the difference in AUROC comparing WGBS and Methyl Array showing that a score derived from the methylation array data generally performs better than one derived from WGBS data.

**Supplemental Figure S5.**
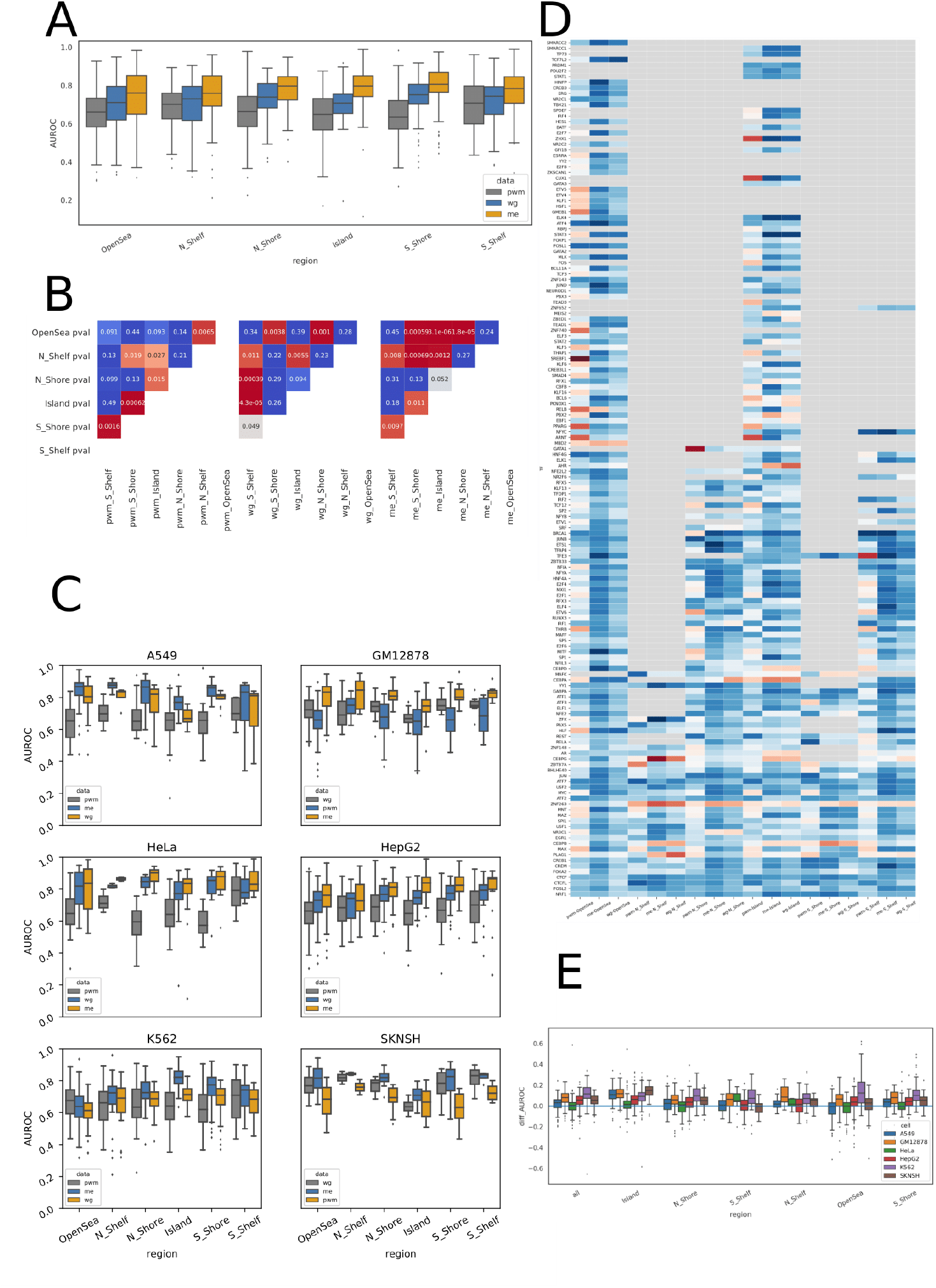
Detailed Analysis based on genomic region. **(A)** Same plot as shown in **Figure 2B**. **(B)** Statistical analyses comparing the distribution of AUROC scores for each pair of regions when scoring motif locations using PWM, WGBS, or methylation array. Shades of red indicate significant differences based on a T-test and associated p-value. **(C)** Cell line specific versions of panel (A). **(D)** Heatmap of specific TF performance averaged across cell lines per gene body proximity. Blue indicates an AUC>0.5 while red indicates AUC<0.5. **(E)** Differential AUROC between methylation array and WGBS indicating that scoring based on methylation array generally performs better than scoring based on the WGBS data.

**Supplemental Figure S6.**
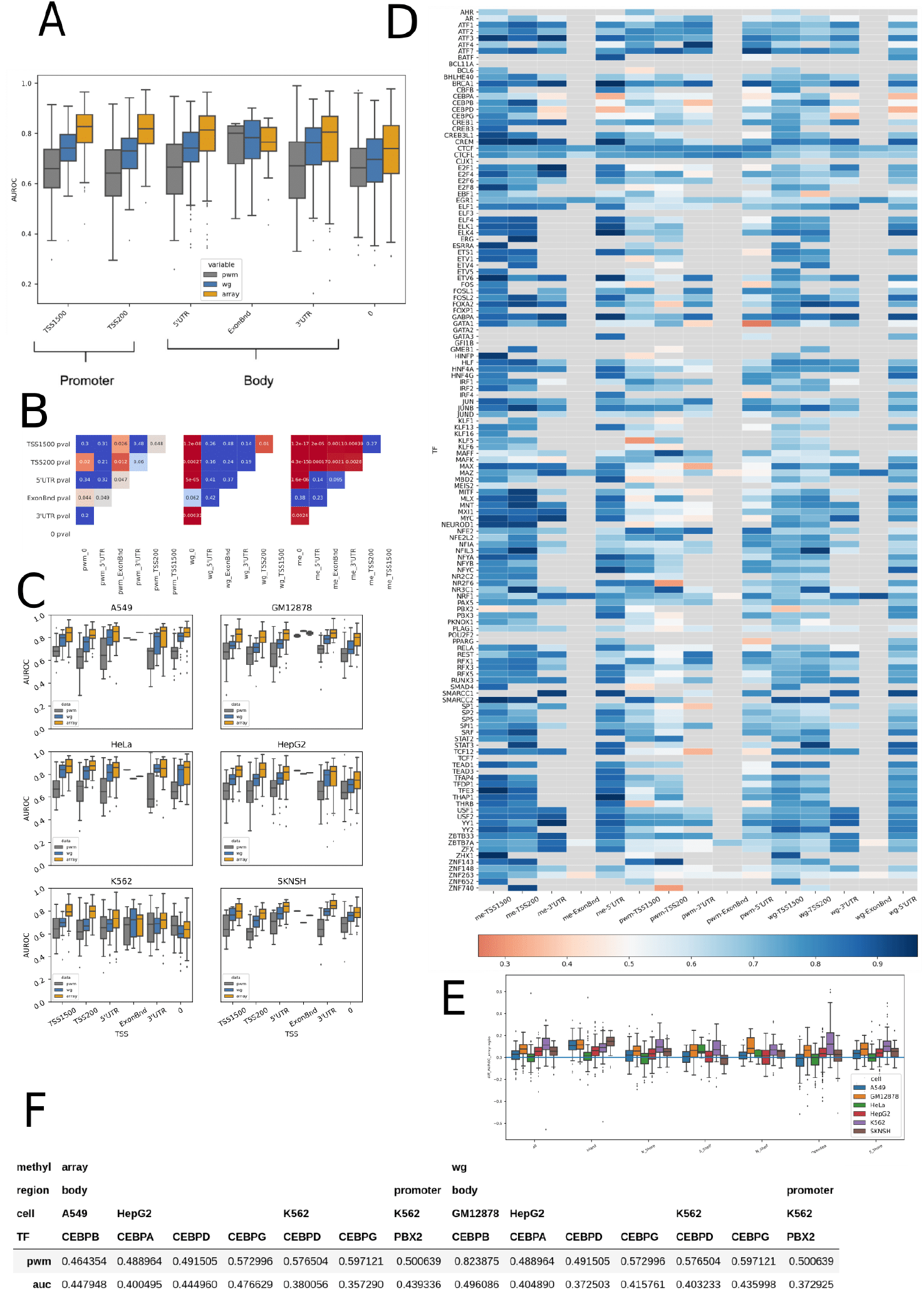
Detailed Analysis based on TSS annotations. **(A)** Same plot as shown in **Figure 2C**. **(B)** Statistical analyses comparing the distribution of AUROC scores for each pair of TSS annotations when scoring motif locations using PWM, WGBS or methylation array. Shades of red indicate significant differences based on a T-test and associated p-value. **(C)** Cell line specific versions of panel (A). **(D)** Heatmap of specific TF performance averaged across cell lines per TSS annotation. Blue indicates an AUC>0.5 while red indicates AUC<0.5. **(E)** Differential AUROC between methylation array and WGBS indicating that scoring based on methylation array generally performs better than scoring based on WGBS data. **(F)** Transcription factors whose context- and cell line specific binding is better predicted using the PWM compared to methylation data and whose methylation-based predictive performance has an AUROC <0.5. Many of these are CEBP family members.

**Supplemental Figure S7.**
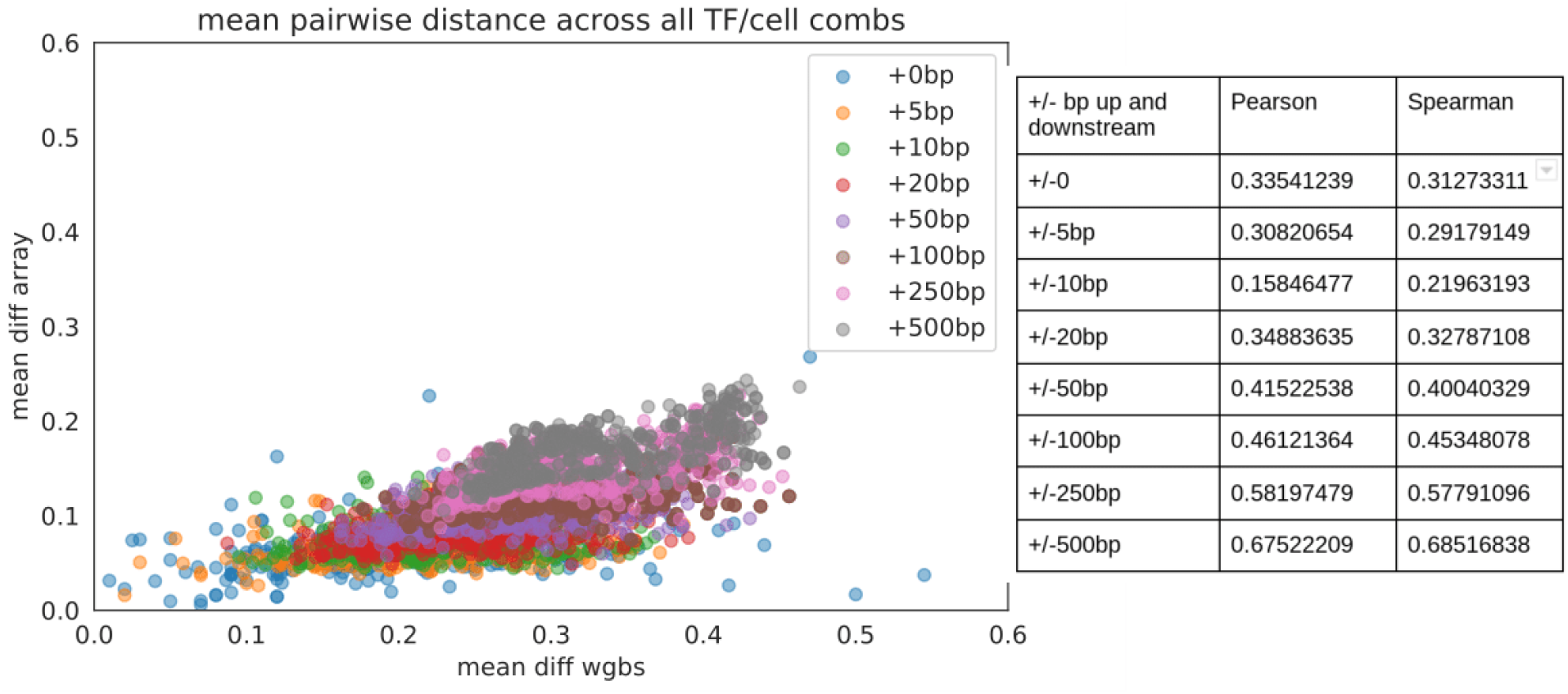
Mean pairwise difference across the multiple CpG methylation values used to score motif locations based on either WGBS or methylation array (each dot is a TF and cell line combination). This plot shows that as the window size is increased the mean pairwise distance between the multiple CpGs used to score motif locations also increases (although this is less so in the array compared to the WGBS) but at the same time becomes more correlated between the two technologies. The accompanying table displays the Pearson and Spearman correlation across TF and cell line values for each window size.

